# Engineered ketocarotenoid biosynthesis in the polyextremophilic red microalga *Cyanidioschyzon merolae* 10D

**DOI:** 10.1101/2023.02.27.530181

**Authors:** Mark Seger, Fakhriyya Mammadova, Melany Villegas-Valencia, Bárbara Bastos de Freitas, Clarissa Chang, Iona Isachsen, Haley Hemstreet, Fatimah Abualsaud, Malia Boring, Peter J. Lammers, Kyle J. Lauersen

**Author notes:** Arizona State University, 7418 Innovation Way South, Mesa, AZ 85212, United States. King Abdullah University of Science and Technology (KAUST), Thuwal 23955-6900, Kingdom of Saudi Arabia.

## Abstract

The polyextremophilic Cyanidiales are eukaryotic red microalgae with promising biotechnological properties arising from their low pH and elevated temperature requirements which can minimize culture contamination at scale. *Cyanidioschyzon merolae* 10D is a cell wall deficient species with a fully sequenced genome that is amenable to nuclear transgene integration by targeted homologous recombination. *C. merolae* maintains a minimal carotenoid profile and here, we sought to determine its capacity for ketocarotenoid accumulation mediated by heterologous expression of a green algal β-carotene ketolase (BKT) and hydroxylase (CHYB). To achieve this, a synthetic transgene expression cassette system was built to integrate and express *Chlamydomonas reinhardtii* (*Cr*) sourced enzymes by fusing native *C. merolae* transcription, translation and chloroplast targeting signals to codon-optimized coding sequences. Chloramphenicol resistance was used to select for the integration of synthetic linear DNAs into a neutral site within the host genome. *Cr*BKT expression caused accumulation of canthaxanthin and adonirubin as major carotenoids while co-expression of *Cr*BKT with *Cr*CHYB generated astaxanthin as the major carotenoid in *C. merolae*. Unlike green algae and plants, ketocarotenoid accumulation in *C. merolae* did not reduce total carotenoid contents, but chlorophyll a reduction was observed. Light intensity affected global ratios of all pigments but not individual pigment compositions and phycocyanin contents were not markedly different between parental strain and transformants. Continuous illumination was found to encourage biomass accumulation and all strains could be cultivated in simulated summer conditions from two different extreme desert environments. Our findings present the first example of carotenoid metabolic engineering in a red eukaryotic microalga and open the possibility for use of *C. merolae* 10D for simultaneous production of phycocyanin and ketocarotenoid pigments.

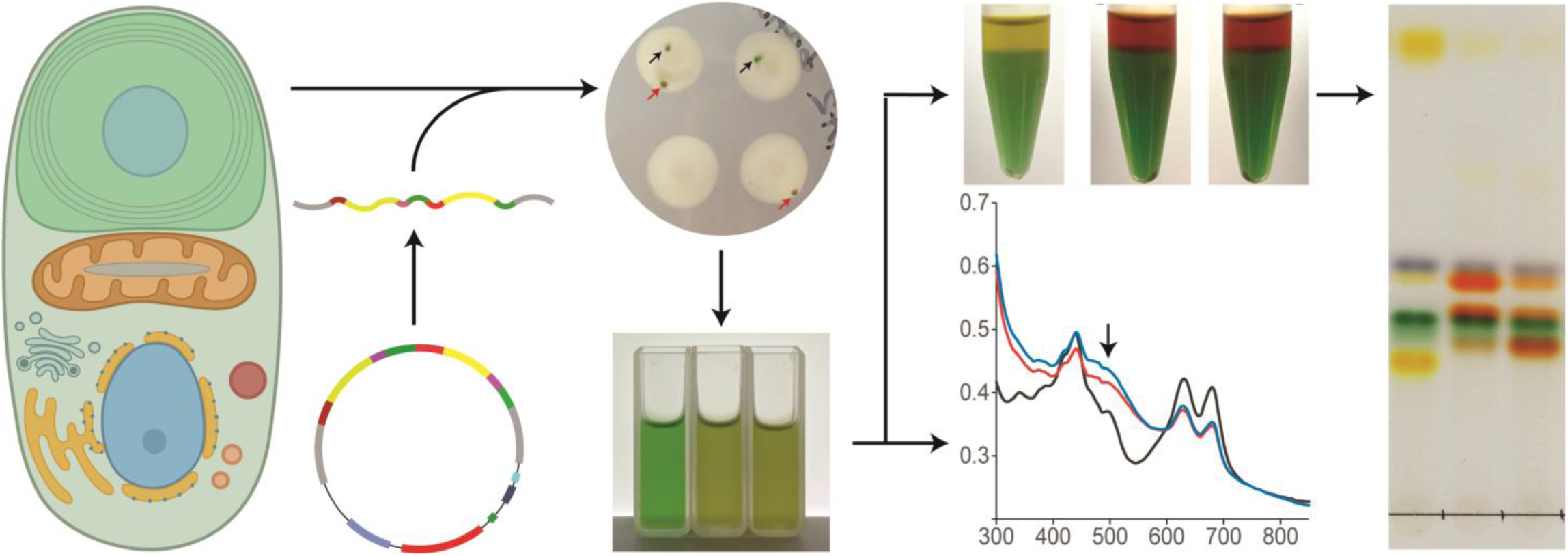

## 1. Introduction

Microalgae are diverse photosynthetic organisms which can be found across the globe in almost every environment, having evolved the capacity for growth on carbon dioxide as a carbon source and the use of (sun)light for energy. Of the many extreme global environments colonized by algae, acidic hot-springs present one of the harshest. Nevertheless, red microalgae from the Class Cyanidiales thrive in water, soil and endolithic environments associated with these hot-springs at temperatures up to 56 °C and pH levels as low as 0.5 (Gross, 2000). The Cyanidiophyceae typically represent the only photosynthetic eukaryotic organisms found tolerating these extreme environments. *Cyanidioscyzon merolae* 10D was isolated from volcanic fields near Naples, Italy (Matsuzaki et al., 2004). It is an obligate photoautotroph with a small genome, one of the first telomere-telomere (∼16 Mbp) complete genome sequences of any model species (Nozaki et al., 2007). Robust tools for genetic manipulation have been developed enabling precise homologous recombination (HR) directed by 200-500 bp targeting sequences (Fujiwara et al., 2017; Takemura et al., 2019a, 2019b). As a result, *Cyanidioschyzon merolae* 10D has emerged as the simplest eukaryotic model cell system with a growing number of useful engineered traits (Miyagishima and Tanaka, 2021), These include the introduction of a cyanobacterial acyl-ACP reductase that resulted in increased triacylglycerol accumulation without growth inhibition (Sumiya et al., 2015) and the incorporation of a Galdieria sulphuraria sugar transporter that enabled heterotrophic growth on glucose (Fujiwara et al., 2019).

The focus of this study is the modification of native carotenoid pigment biosynthesis in *C. merolae* 10D. Ironically, the red microalgae are blue-green in color like cyanobacteria as they share the trait of phycocyanin use as a light-harvesting photopigment and only contain chlorophyl a. *C. merolae* 10D has a minimal carotenoid profile lacking alpha-carotene and lutein, it accumulates β-carotene and zeaxanthin as its terminal carotenoids and completely lacks violaxanthin and neoxanthin (Figure 1) (Cunningham et al., 2007). The capacity for HR transgene integration into its nuclear genome, minimal intron content, and general ease of handling make *C. merolae* 10D an exciting candidate for green (red) synthetic biology and metabolic engineering investigations (Lang et al., 2020; Pancha et al., 2021). Its extreme growth requirements also allow *C. merolae* to be cultivated with minimal risk of contamination and could be a promising host for industrial-scale algal waste-stream conversion processes (Delanka-Pedige et al., 2019; Selvaratnam et al., 2022). In addition, Cyanidiales phycocyanin is more thermostable than that currently sourced from *Arthrospira platensis* (Spirulina) and is a potentially valuable co-product which can be a co-product from engineered cell biomass (Rahman et al., 2017).

**Figure 1.**
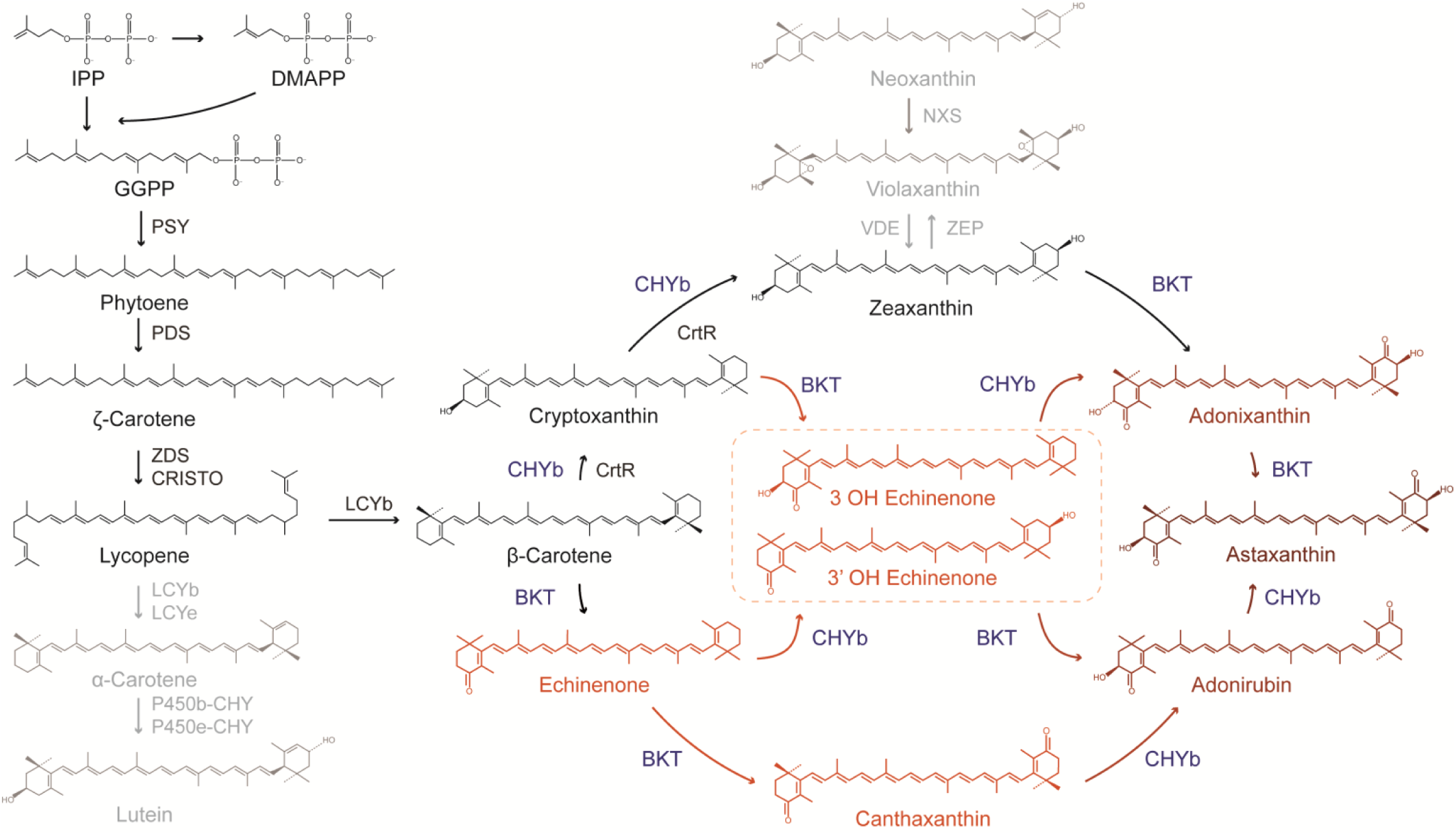
Carotenoid pathway of *C. merolae* and its extension to ketocarotenoid biosynthesis. *C. merolae* 10D lacks the α-carotene branch of carotenoid biosynthesis and accumulates only the terminal xanthophyll zeaxanthin but not violaxanthin or neoxanthin. Pathways not found in C. merolae are shown in light grey. Native carotenoid pathway enzymes are shown in black, heterologous BKT and CHYB are shown in blue. BKT acts to add ketone groups to the terminal carotenoid rings, while CHYB hydroxylates them, yielding several intermediates in the production of astaxanthin. Ketocarotenoids and intermediates are shown in orange and red. Chemical abbreviations: IPP, DMAPP, and GGPP – isopentyl, dimethylallyl, and geranylgeranyl pyrophosphate. Gene names: PSY – phytoene synthase, PDS – phytoene desaturase, ZDS/CHRISTO – ?-carotene desaturase/carotene isomerase, LCYb – lycopene β-cyclase, LCYe – lycopene ε-cyclase, P450b/e-CHY – P450-carotene hydroxylases, CrtR – β-carotene hydroxylase (cyanobacterial), VDE – violaxanthin de-epoxidase, ZEP zeaxanthin epoxidase, NXS – neoxanthin synthase.

Recently, advances in transgene design opened metabolic engineering in the green model microalga *Chlamydomonas reinhardtii*, in which native carotenoid profiles have been modified to produce the ketocarotenoids canthaxanthin and astaxanthin (Amendola et al., 2023; Lauersen, 2019; Perozeni et al., 2020). Both ketocarotenoids have value for their high antioxidant properties, application as food coloring, as well as pharmacological uses (Ambati et al., 2014). Bulk production of ketocarotenoid pigments would help drive the transition to non-toxic, natural textile dyes (Shabbir et al., 2018). Carotenoid modification in the green alga was achieved by overexpression of its native β-carotene ketolase (*Cr*BKT) and hydroxylase (*Cr*CHYB) in vegetative green cells where they are not naturally expressed (Amendola et al., 2023; Perozeni et al., 2020). Overexpression *Cr*BKT resulted in color changes of the green algal cells to brown due to global changes in pigment composition - the accumulation of orange-red ketocarotenoids and both chlorophyll a and b (Cazzaniga et al., 2022; Perozeni et al., 2020). In *C. reinhardtii, Cr*BKT expression alone generates intermediate ketolated carotenoids from native β-carotene, zeaxanthin substates, and partially hydroxylated carotenoids to form canthaxanthin, intermediates, and small amounts of astaxanthin. Recently, it was shown that the hydroxylation of these to astaxanthin was enhanced by co-overexpression of *Cr*CHYB in *C. reinhardtii* (Amendola et al., 2023).

Here, the capacity for carotenoid engineering in the model red microalga *C. merolae* 10D was investigated. As part of this work, a completely synthetic plasmid toolkit was built and tested, with domestication of transcriptional elements, targeting peptides, and protein tags optimized for expression of target transgenes from either one- or two-gene cassette(s) from the nuclear genome of *C. merolae* 10D. The green algal BKT and CHYB were optimized for the red algal nuclear genome context and expressed in fusion protein constructs from these plasmids after genomic integration in the intergenic region found in the 184-185C locus of *C. merolae* 10D chromosome 4. Transformants with confirmed HR integration of transgenes exhibited expression of each target product and colorimetric changes to culture pigmentation caused by ketocarotenoid accumulation which were visible by eye. The effects on cellular pigments were quantified and documented. Unlike in green algae, total carotenoids were not reduced in *C. merolae* 10D when ketocarotenoids were produced and these pigments did not affect cellular phycocyanin titers. Growth behaviors were investigated in optimal and modeled extreme desert environments using programmed bioreactors to show the potential for scaled cultivation concepts with engineered keto-carotenoid producing *C. merolae* 10D. Our results indicate that the polyextremophile is readily amenable to genetic manipulation, its carotenoid profile can be modified to generate ketocarotenoids, and future bioprocesses could harvest these separately from water-soluble phycocyanin. Here we started with a red alga which looks cyan and used green algal carotenoid biosynthetic genes to turn make it red-brown while not impacting its blue pigment composition. Our findings encourage further investigations of metabolic engineering with this promising eukaryotic photosynthetic cyan-cell chassis.

## Materials and Methods

### 2.1 Algae culture

The strain of *C. merolae* 10D (wildtype; NIES-3377) was obtained from the National Institute of Environmental Studies’ microbial culture collection in Japan. The culture was immediately plated on corn starch beds (Ohnuma et al., 2008) and single colonies were isolated, scaled, and verified as mono-algal cultures using microscopy and PCR. These cultures along with its transgenic lines were maintained in MA2 medium (Kuroiwa et al., 2017), which consists of 40 mM (NH_4_)_2_SO_4_, 8 mM KH_2_PO_4_, 4 mM MgSO_4_, 1 mM CaCl_2_, 100 μM FeCl_3_, 72 μM EDTA-2Na, 16 μM MnCl_2_, 2.8 μM ZnCL_2_, 7.2 μM NaMoO_4_, 1.3 μM CuCl_2_, and 0.7 μM CoCl_2_. The pH was adjusted to 2.3 with H_2_SO_4_. For long term preservation, verified cultures were cryopreserved in 13.5% DMSO using Quick-freezing containers (Mr. Frostys™, Thermo Sci.) at -80 °C. Working stocks of cultures were maintained on corn starch beds on MA2 Gellan gum plates and in TC flasks (CELLTREAT^®^; USA) with constant agitation under continuous illumination (200 μmol m^-2^ s^-1^) at 40 °C in Percival incubators (Percival Scientific; USA) supplemented with 3% CO_2_ mixed in air.

### 2.2 *In silico* genetic designs

Eight transformation plasmids were designed and constructed as follows to integrate the selectable marker (chloramphenicol acetyltransferase (CAT)) and transgenes (mVenus (YFP), β-carotene ketolase (BKT), and β-carotene hydroxylase (CHYB)) cassettes into the intergenic region between the nuclear glycogen phosphorylase (CDM184C) and TATA-box binding protein-associated factor 13 (CMD185C) genes via homologous recombination. The origin, sequences, primers, and references for the genetic control elements used in our *in silico* design process are listed in Supplemental Data S1 and S2. Endogenous sequences (regulatory elements, transit peptides, and homology arms) were extracted from the reference genome of *C. merolae* 10D (Fujiwara et al., 2019, 2017, 2013; Moriyama et al., 2014). The CAT, YFP, NOS terminator, and BKT/CHYB sequences, derived from *S. aureus, A. victoria, A. tumefaciens*, and *C. reinhardtii (Cr)* (respectively), were taken from the literature/NCBI database (Amendola et al., 2023; Hopp et al., n.d.; Kremers et al., 2006; Perozeni et al., 2020; Schmidt et al., 2007; Sumiya et al., 2014; Zienkiewicz et al., 2017). Codon optimization of coding sequences (CDS’s), along with removal of unwanted restriction sites, was carried out using Geneious Prime (v. 2023.0.1; Biomatters Lt., New Zealand) and *C. merolae*’s codon usage table found in the Kasusa database (https://www.kazusa.or.jp/codon/cgi-bin/showcodon.cgi?species=280699). Restriction enzyme recognition sequences for the enzymes listed at the top of Fig. 2A were systematically removed from all sequences used in our one- and two-cassette constructs.

**Figure 2.**
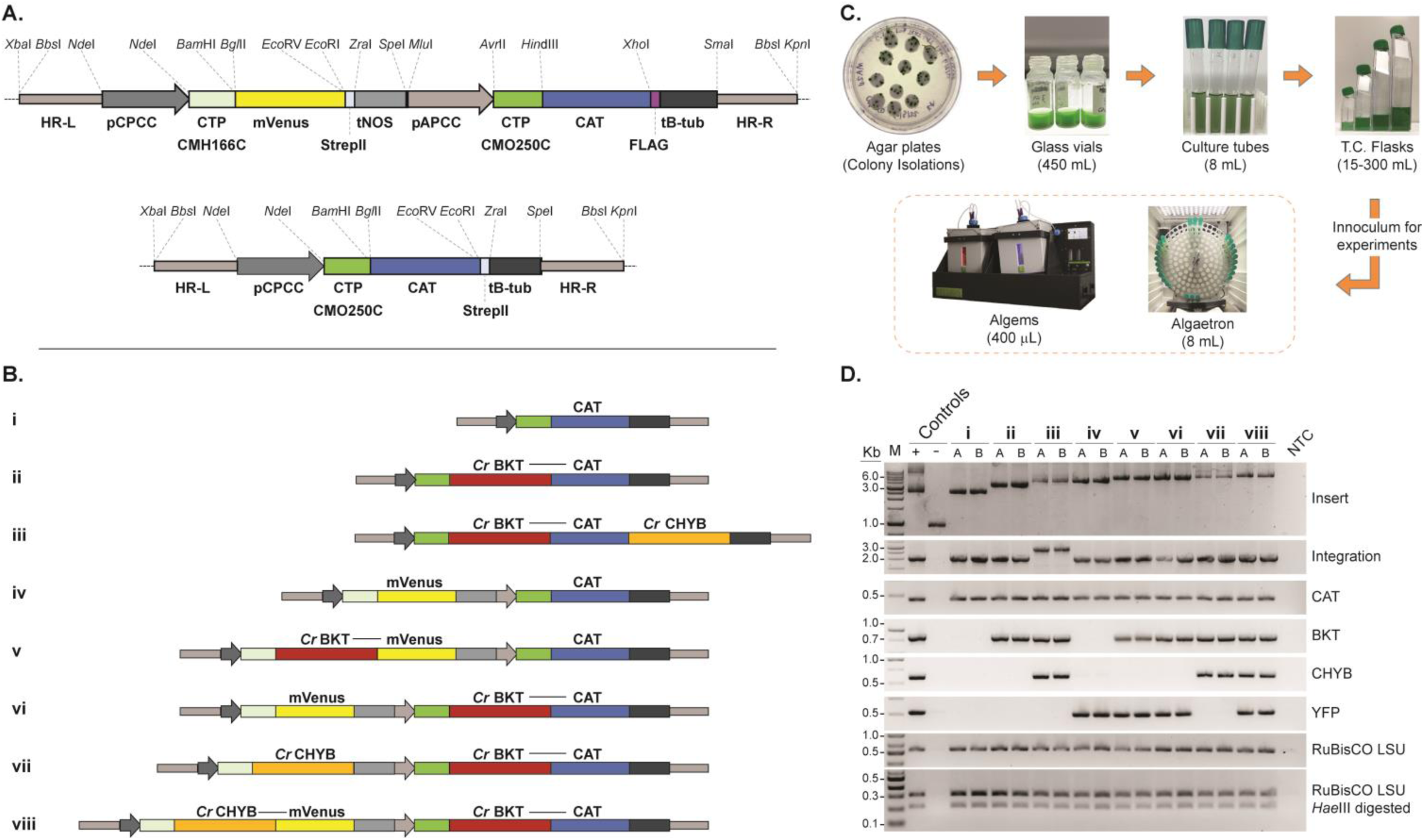
Plasmid design, culturing systems and transgene integration. A – Synthetic plasmids were designed *in silico* and constructed *de novo* for integration of transgenes into the 184C-185C locus (HR-L and -R) on *C. merolae* chromosome 4. Two template plasmids were synthesized: a two-cassette (upper) and a single cassette (lower), both with chloramphenicol (CAT) resistance marker as a selection/fusion partner. Expression elements and gene fragments are separated by non-redundant restriction endonuclease sites as illustrated. pCPCC – phycocyanin-associated rod linker protein promoter, CTP CMH166C – DNA Gyrase B chloroplast targeting peptide, mVenus – yellow fluorescent protein reporter, StrepII – C-terminal peptide tag with stop codon, tNOS – nopaline synthase terminator,pAPCC – allophycocyanin-associated rod linker protein promoter, CTP CMO250C – allophycocyanin-associated rod linker protein chloroplast targeting peptide, FLAG – peptide tag with stop codon, tB-tub – *C. merolae* β-tubulin terminator CMN263C. B – *C. reinhardtii* β-carotene ketolase (*Cr*BKT) and β-carotene hydroxylase (*Cr*CHYB) transgenes were codon optimized for *C. merolae* nuclear genome expression based on amino acid sequences and native targeting peptide removal and subcloned into either of the above two plasmids as illustrated for expression as either target-mVenus or -CAT fusion proteins. C – transformation of *C. merolae*, recovery of colonies in starch spots on chloramphenicol selection, and seed train for experiments. D – polymerase chain reaction confirmation of plasmid integration at the 184-185C neutral locus, presence of transgenes, and unialgal status (RuBusCO *Hae*III digestion). Information on primers and PCR assays found in Supplemental Figure S1 and Data S2.

Thus, intermediate constructs used to create the eight constructs used here are available to speed future designs. For regulatory elements and homology arms, silent single point mutations (SPMs) were introduced manually in sequences to remove unwanted restriction sites. Modified promoter and terminator sequences were analyzed and compared to original sequences using Softberry Nsite(M)-PL (www.softberry.com) and Geneious DNA-fold (v. 2023.0.1; Biomatters Lt., New Zealand) to ensure conserved regulatory motifs and secondary structures, respectively, were not altered. All SPMs were documented and are indicated in sequences as lower-case bases (Supplemental Data S1). *In silico* assembly and *de novo* synthesis of transformation plasmids using pBluesript II KS (+) (Stratagene, USA) as the backbone vector was done in the Snapgene (software v. 6.4; www.snapgene.com) and using GenScript services (GenScript Inc., USA), respectively (Figure 2). All plasmids were transformed into chemically competent *E. coli* JM109 cells and plasmids were extracted using ZymoPURE II midiprep kits (Zymo Research group, California).

### 2.3 C. *merolae* 10D transformation

To prepare linear DNA fragments for transformation, PCR was performed using primer set 1 (detailed in Supplemental Figure S1 and Data S2) and plasmid DNA. The resulting products were then purified by ethanol precipitation. PEG-mediated transformation of *C. merolae* 10D was carried out using four micrograms of linear DNA as previously described (Fujiwara et al., 2021, 2013) with some modifications. Transfected cells were transferred into 8.0 mL of MA2 media in 20 mL culture tubes and allowed to recover while rotating (∼80 rpm) in the outer rim of a tissue culture roller drum (New Brunswick; model TC-7; Eppendorf, USA) housed in an Algatron^®^ incubator (Photon Systems Instruments, Czech Republic) supplemented with 3% CO_2_ in air with continuous illumination (100 μmol m^-2^ s^-1^) at 40 °C for two days. Cells were subsequently collected by centrifugation, supernatant discarded, and cells resuspended in MA2 (∼400 μL).

Cell suspensions were then serial diluted in MA2 and 200 μL of the dilutions were amended to equal volume of 40% corn starch with chloramphenicol (“Cm” 300 μg/mL). Approximately 20 μL aliquots of cornstarch slurry with cells were spotted on MA2 agar (0.5%) plates (60 × 90 mm) with Cm [150 μg/mL]. Plates, with approximately 18-20 inoculated cornstarch beds, were incubated in humidified CO_2_ chambers under the same conditions described above until colony formation. At which point colonies were isolated, transferred into 400 μL of MA2 with Cm [150 μg/mL] in 2.0 mL glass vials, and then allowed to grow for ∼7-10 days. Isolates were screened using a colony PCR method with primer set 2 (Supplemental Figure S1 and Data S2) to test for integration of our cassettes into the targeted neutral site. Positive transformants were then scaled up as shown in Figure 2C and characterized via PCR, flow cytometry, fluorescent microscopy, UV–vis spectrophotometry, thin layer chromatography (TLC), and high-performance liquid chromatography (HPLC). A subculture from each was cryopreserved in DMSO for long term storage (as described above).

### 2.4 DNA extractions and PCR assays

Cultures were harvested by centrifugation (5 min at 14,000xg) and total genomic DNA was extracted from algal cell pellets (∼50-100 mg) with a Zymo Quick-DNA fungal/bacterial extraction kit (Zymo Research group, USA) according to the manufacturer’s protocol. DNA extracts were quantified using a NanoDrop One spectrophotometer (Thermo Fisher Scientific, USA). The high fidelity PrimeSTAR GXL DNA Polymerase (Takara Bio Inc., Japan) and the Hot start GoTaq polymerase (Promega Corporation, USA) were used for PCR according to the manufacturer’s protocols. The former was specifically used with primer set 1 to amplify the insert DNA (HR-L to HR-R) for transfection and to screen transformants for presence of the insert at the target neutral site. All primers used to screen cultures were synthesized by IDT (Integrated DNA Technologies Inc., San Diego) and primer sequences along with PCR conditions and relative primer annealing sites are shown in Supplemental Data S2 and Figure S1, respectively.

### 2.5 UV–vis spectrophotometry

A HACH DR5000 UV–Vis spectrophotometer was used to monitor culture growth by measuring the optical density at 750 nm and to analyze pigment extracts, unless otherwise stated. *In vitro* spectral profiles of wild type and transformed cells were obtained using a SpectraMax i3 plate reader (Molecular Devices, CA, USA) across a range of wavelengths spanning from 300-850 nm.

### 2.6 Epifluorescence microscopy

Cells were visualized and imaged with 100X objective lens and immersion oil using an Olympus BX51 fluorescence microscope equipped with a Canon EOS RP DSLR camera. Fluorescence microscopy was performed on transformants specifically expressing the mVenus (YFP) transgene to verify localization of YFP in the chloroplast and evaluate cassettes with YFP fusions. Two different excitation filters were used for detecting pigment and YFP fluorescence: U-MWG2 and FITC-3540B-OMF, respectively.

### 2.7 Flow cytometry

Flow cytometric analyses of wild type and YFP transformant cells was performed using a Guava^®^ easyCyte™ HT BGV flow cytometer (Luminex Corporation, Austin, TX, USA) equipped with a blue (488 nm) laser; which was used to measure size (forward scatter), granularity (side scatter), chlorophyll fluorescence (692/40 nm) and YFP fluorescence (575/25 nm). All samples were normalized to 0.01 OD_750_ (∼350-450 cells μL^-1^) and a total of 10,000 events were recorded per sample. Data acquisition and analysis was done using GuavaSoft v. 3.4 software (InCyte; Luminex Corporation).

For the Algem photobioreactor growth experiment, the cell densities were measured using an Invitrogen Attune NxT flow cytometer (Thermo Fisher Scientific, UK) equipped with a Cytkick microtiter plate autosampler unit as recently described (de Freitas et al., 2023). Each sample was diluted 1:100 with 0.9% NaCl solution and loaded into a 96-well microtiter plate in technical triplicates, the cell density was measured from this plate using the autosampler. Samples were mixed three times immediately before analysis, and the first 25 μL of the sample was discarded to ensure a stable cell flow rate during measurement. For the data acquisition, 50 μL from each well was analyzed.

### 2.8 Biomass determination

For 20 mL culture tube growth experiment, Ash-free dry weights were determined using OD_750_ values and an OD_750_ to AFDW correlation coefficient, which was determined for each transformant prior to the experiment and found to be the same for all strains: AFDW (g/L) = 0.27 * (OD_750_ nm). This correlation coefficient was determined as previously described (Dandamudi et al., 2021). For Algem photobioreactor growth experiments, biomass was measured by vacuum filtration of 4 mL of each test on pre-weighted filters (0.45μm). The algal cells were dried at 60 °C for 24h in petri dishes, then allowed to cool before weighing the filter with the biomass. All measurements consisted of technical and biological triplicates.

### 2.9 Pigment extraction and analysis

All extractions and analyses of pigments were carried out in dark or dim light to avoid photodegradation. For phycocyanin extraction, 4.5 mg of freeze-dried biomass was added into 1.5 mL 0.1M phosphate buffer (pH 7.0) and subjected to bead beating (Bullet Blender^®^ STORM 24, Next Advance, USA) using a mix of 0.15 mm and 0.5 mm zirconium oxide beads at the highest speed for 5 min. The supernatant was recovered by centrifugation at 12,000xg for 5 min, and the pellet was re-extracted under the same conditions. Both supernatants were combined and analysed spectrophotometrically.

The extraction of carotenoids and chlorophyll *a* was performed using 10 mg of freeze-dried biomass added to 800 μL of acetone containing 0.1% (w/v) butylated hydroxytoluene to prevent carotenoid oxidation. The mixture was homogenized via bead beating as described above. The supernatant was collected after centrifugation at 12,000xg for 3 min, and the remaining pellet was subjected to three additional extractions using 600 μL of acetone until the supernatant became colorless. All the supernatants were pooled and evaporated to dryness under a stream of nitrogen.

For carotenoid saponification, dried extracts were resuspended in 300 μL ethyl acetate and treated with 300 μL 5% (w/v) methanolic KOH under constant shaking at room temperature for 2 hr. To stop the reaction 100 μL of 10% (w/v) NaCl, and 200 μL of deionized water were added to the reaction mixture, and carotenoids were extracted four times with hexane:MTBE (1:1, v/v, 300 μL per extraction) using centrifugation (12,000xg, 1 min) to separate the layers. The organic layers were collected and combined, then evaporated to dryness under a stream of nitrogen. Dried extracts, whether saponified or non-saponified, were dissolved in 1 mL of acetone, filtered using a 0.45 μm nylon filter in preparation for pigment analysis by TLC, UV-Vis spectrophotometry and HPLC.

TLC was used to separate and identify carotenoids. 20 μl aliquots of the pigment extracts and carotenoid standards were spotted on pre-coated silica gel 20×20 cm TLC plates (company info) and eluted with a mobile phase of hexane:acetone (7:3, v/v). The concentrations of phycocyanin, chlorophyll *a* and total carotenoids were determined spectrophotometrically. The absorbance of phycocyanin extracts was measured at 620 and 652 nm, and the concentration of phycocyanin was calculated using previously published equations (Bennett and Bogorad, 1973). For the assessment of chlorophyll *a* and total carotenoid contents, absorbance of extracts was recorded at 662 and 470 nm, respectively, and the concentrations of chlorophyll *a* and total carotenoids were calculated according to previously published equations. Separation of carotenoids and their quantification were conducted by reverse-phase HPLC (Waters Alliance 2695 Separations Module coupled with a 2996 photodiode array detector) as described in (Amendola et al., 2023; Perozeni et al., 2020). The HPLC system was equipped with a C18 column (Waters Spherisorb ODS2 Column 5 µm, 4.6 mm × 250 mm, Supelco, Inc., Belefonte, PA, USA) and a 15 min gradient of ethyl acetate (0%–100%) in acetonitrile–water–triethylamine (9:1:0.01, v/v/v) was employed at a flow rate of 1 mL/min. Carotenoid peaks were identified by comparing retention times and spectra to carotenoid standards, which were also used to quantify carotenoids using standard curves (Supplemental Data S3).

### 2.10 Growth experiments

#### 2.10.1 Culture tube experiment under different light conditions

Wild type and transformant (ii and viii) inoculates were preadapted at 750 μmol m^-2^ s^-1^ in Algatron^®^ incubators (Photon Systems Instruments, Czech Republic) under the same conditions as described above (with the exception of the light conditions) for 5 days. Biomass was collected by centrifugation and pellets resuspended in fresh MA2 medium (pH 2.3) with a starting density of 0.8 OD_750_. Triplicate sets of 20 mL culture tubes were prepared (8.0 mL working volume) for each test strain for each light condition (750 and 1172 μmol m^-2^ s^-1^). A total of 3 sets were prepared: one for daily growth metrics and the other two for pigment analysis. The culture tubes were arranged in the outer rim of a tissue culture roller drum that was housed in an Algatron^®^ incubator as above, according to their respective light conditions. Water acidified to medium pH was added as need to account for evaporative losses. When sampling daily for growth metrics (≤100 μL), the same volume that was removed for sampling was replaced with medium. Culture density was monitored spectrophotometrically (as described above). The two sacrificial sets of tubes for pigment analysis were collected at different growth phases: one at log phase and the other at stationary phase. Biomass was collected from each culture tube by centrifugation (4,200xg for 10 min) and pellets freeze dried for pigment analysis. Growth metrics and pigments analysis (N = 3x biological and technical replicates) was done as described in the previous sections.

#### 2.10.2 Algem photobioreactor performance benchmarking in modelled environments

*C. merolae* 10D WT and transformant lines ii and viii were first precultured in MA2 liquid medium (pH 2.3) in 125 mL Erlenmeyer flasks with a working volume of 10 mL for 4 d under continuous agitation (100 rpm) and illumination (90 μmol photons m^-2^ s^-1^) in a CO_2_ (4%) incubator at 42 °C. These cultures were then used to inoculate 1 L Algem photobioreactor flasks (working volume 400 mL) with a target density of 3 ×10^6^ cells mL^−1^. To simulate outdoor light and temperature conditions of Thuwal, Saudi Arabia (22.3046N, 39.1022E) and Mesa, Arizona, United States (33.305130N, -111.67300W), environmental conditions were developed based on data sets reported by (de Freitas et al., 2023) and obtained from the AzCATI facility, respectively. Four different growth conditions were used to evaluate the growth performance of each strain: (1) constant light (1500 μmol m^-2^ s^-1^) and temperature (42 °C), (2) 12:12 h light:dark with these same light and temperature conditions, and simulated seasonal environmental conditions of (3) Thuwal and (4) Mesa, with the months of February, May, August, and November representing winter, spring, summer, and autumn (respectively). Samples of 15 mL were collected daily for cell concentration, biomass quantification, and carotenoid analysis as described above. The same volume that was removed for sampling was replaced with sterilized water acidified to medium pH.

## 3. Results and Discussion

The polyextremophile *C. merolae* 10D is restricted to low pH (0.5-5) and temperatures from 35-56 °C (Miyagishima and Tanaka, 2021). It has a simplified natural carotenoid profile which lacks the alpha-branch of carotenoid biosynthesis and has only β-carotene and zeaxanthin as terminal carotenoids (Figure 1, (Cunningham et al., 2007)). In higher plants and green algae, alpha-carotene is converted into lutein, and zeaxanthin is used to create violaxanthin and neoxanthin as part of the photoprotective/photoresponsive xanthophyll cycle (Goss and Jakob, 2010; Latowski et al., 2004). As these pigments are absent in *C. merolae*, it is an interesting species with a simplified carotenoid substrate and biosynthesis enzymatic landscape in which to attempt carotenoid metabolic engineering. *C. merolae* also uses phycocyanin as a light harvesting pigment (Lang et al., 2020; Parys et al., 2021), a different photosystem structure than in green algae and higher plants, opening the question what effects carotenoid modifications would have in this system.

Ketocarotenoids are orange-red pigments that are formed through the ketolation of the terminal rings of β-carotene and zeaxanthin to form a range of intermediates towards canthaxanthin (dual-ketolated β-carotene) and astaxanthin (dual-ketolated and hydroxylated β-carotene) (Figure 1) (Perozeni et al., 2020). Canthaxanthin and astaxanthin are formed in a range of organisms including algae, plants, bacteria and fungi (Seybold and Goodwin, 1959; Wan et al., 2021; Zhang et al., 2020). These pigments have various applications from food colorants, aquaculture feed enhancements, medicinal treatment of skin diseases, as specialty chemical conjugants, and are considered powerful antioxidants (Ambati et al., 2014). Recent reports have shown that it is possible to leverage gene redesign and synthetic overexpression of the native β-carotene ketolase (*Cr*BKT) and hydroxylase (*Cr*CHYB) of the green microalga *C. reinhardtii* to produce canthaxanthin, intermediate ketocarotenoids, and astaxanthin in this photosynthetic microbe (Amendola et al., 2023; Cazzaniga et al., 2022; Perozeni et al., 2020). The BKT adds ketone groups to the terminal rings of both zeaxanthin and β-carotene, while CHYB adds hydroxyl-groups to β-carotene (Figure 1) (Amendola et al., 2023). As both β-carotene and zeaxanthin are the terminal carotenoids within *C. merolae* and its growth conditions minimize risk of contaminating organisms, we reasoned it could be an efficient cell chassis for metabolic engineering and biotechnological ketocarotenoid production.

### Synthetic transgene expression cassette design and transformation

Recent reports indicated the possibility of nuclear transformation and efficient transgene integration by homologous recombination (HR) in *C. merolae* (Fujiwara et al., 2021, 2019, 2017, 2013; Minoda et al., 2004; Takemura et al., 2019a, 2018). Here, it was investigated whether a synthetic-biology strategy could be used to enable heterologous expression of the green algal ketocarotenoid-biosynthesis enzymes in *C. merolae*. Promoter, terminator and plastid targeting signals (Miyagishima and Tanaka, 2021) were used to drive expression of *C. merolae* codon optimized sequences coding for *Cr*BKT and *Cr*CHYB *in silico* (Figure 2A) and the expression cassettes commercially synthesized *de novo*. The expression cassettes were designed to be modular, with each element separated by unique restriction endonuclease sites and a previously demonstrated target for HR was chosen, the 184-185C locus found on *C. merolae* 10D chromosome 4 (Fujiwara et al., 2017). Coding sequences for each target transgene were optimized for the *C. merolae* codon usage bias before synthesis and selection was achieved with a codon optimized chloramphenicol resistance (CAT) marker. Plasmids were built to express *Cr*BKT and *Cr*CHYB in various fusion constructs to either the CAT resistance marker or yellow fluorescent protein (mVenus, YFP) in different combinations of gene cassettes (Figure 2B). Full, annotated sequences of all plasmids are provided in Supplemental Data S4.

To enable expression of the *Cr*BKT and *Cr*CHYB, different genetic fusion constructs were used to allow selection for expression with either antibiotic resistance or visually through fluorescence screening (Figure 2B). Plasmid i was designed to express the chloramphenicol resistance marker (CAT) and localize it to the algal plastid with a targeting peptide of a native protein. Transformants generated with this act as controls for other constructs. Similarly, construct iv serves as a control for the fluorescent reporter mVenus (YFP), which was also targeted to the algal plastid through a separate targeting peptide than the CAT resistance marker (Supplemental Figure S2). *Cr*BKT has been shown to be a highly active enzyme in the production of ketocarotenoids and is effective in direct fusion to the spectinomycin resistance marker in *C. reinhardtii* (Amendola et al., 2023; Cazzaniga et al., 2022). We emulated this strategy of selection marker fusion to the *Cr*BKT here (constructs ii, iii, vi, vii, viii) with CAT as this selection marker functions to yield resistance colonies in *C. merolae* 10D and also functions when localized in the algal plastid where carotenoid biosynthesis occurs (Minoda et al., 2004). Fusion to a reporter protein can also increase the half-life of target recombinant proteins in cells and improve overall to target product yields in metabolic engineering efforts (Cheah et al., 2022). This strategy has been effective in overcoming nuclear transgene expression limitations in green algae, and was shown to be the most effective strategy for *Cr*BKT fusion in its original report (Lauersen, 2019; Perozeni et al., 2020). Therefore, construct v was designed to express *Cr*BKT in fusion with YFP to determine if it was more effective than CAT fusion. *Cr*CHYB was shown to express well in *C. reinhardtii* where it catalyzes hydroxylation of ketocarotenoids to astaxanthin (Amendola et al., 2023). Here, we chose to attempt its expression alone (vii), in fusion with YFP (viii), or in longer fusion to the C-terminus of *Cr*BKT-CAT (iii). Each was investigated to determine whether binary cassettes of larger sizes could be integrated into the genome by HR, and whether different efficacy in astaxanthin biosynthesis could be achieved with different fusion orientations (Figure 2B).

Synthetically designed plasmids were used as templates for PCR to generate linear DNA fragments used in PEG-mediated transformation of *C. merolae*. Colonies of *C. merolae* 10D resistant to chloramphenicol could be readily achieved in starch beds following reported protocols (Minoda et al., 2004) for every construct designed in this work (Figure 2C). Colonies were isolated by picking and grown in 400 μL MA2 liquid medium in standing glass vials prior to further analysis. For each plasmid construct, several dozen colonies were selected and checked for integration by PCR using primers listed in Supplemental Data S2. Representative clones were used to show profiles of PCR products indicating genomic integration markers (Figure 2D) and representatives from each transformant pool used in carotenoid analysis. Expression success is described in the following section in relation to effects on carotenoid biosynthesis.

### β-carotene ketolase and hydroxylase generate ketocarotenoids in *C. merolae* 10D

All carotenoid modifying enzymes were successfully expressed from our synthetic transgene constructs in *C. merolae* 10D and caused changes to the native carotenoid profiles in each strain (Figure 3). This effect was visible already in cultures to the naked eye (Figure 3A) and was confirmed by spectrophotometric scans (Figure 3B) similar to those previously reported for *Cr*BKT and *Cr*CHYB expression in Chlamydomonas (Amendola et al., 2023; Perozeni et al., 2020). It was observed here that all *C. merolae* 10D transformants with *Cr*BKT or *Cr*BKT+*Cr*CHYB expression exhibited a visible color change relative to the parental strain (Figure 3A). Absorbance measurements revealed a shoulder at ∼500 nm, a phenotype previously reported in organisms accumulating ketocarotenoids (Figure 3B).

**Figure 3.**
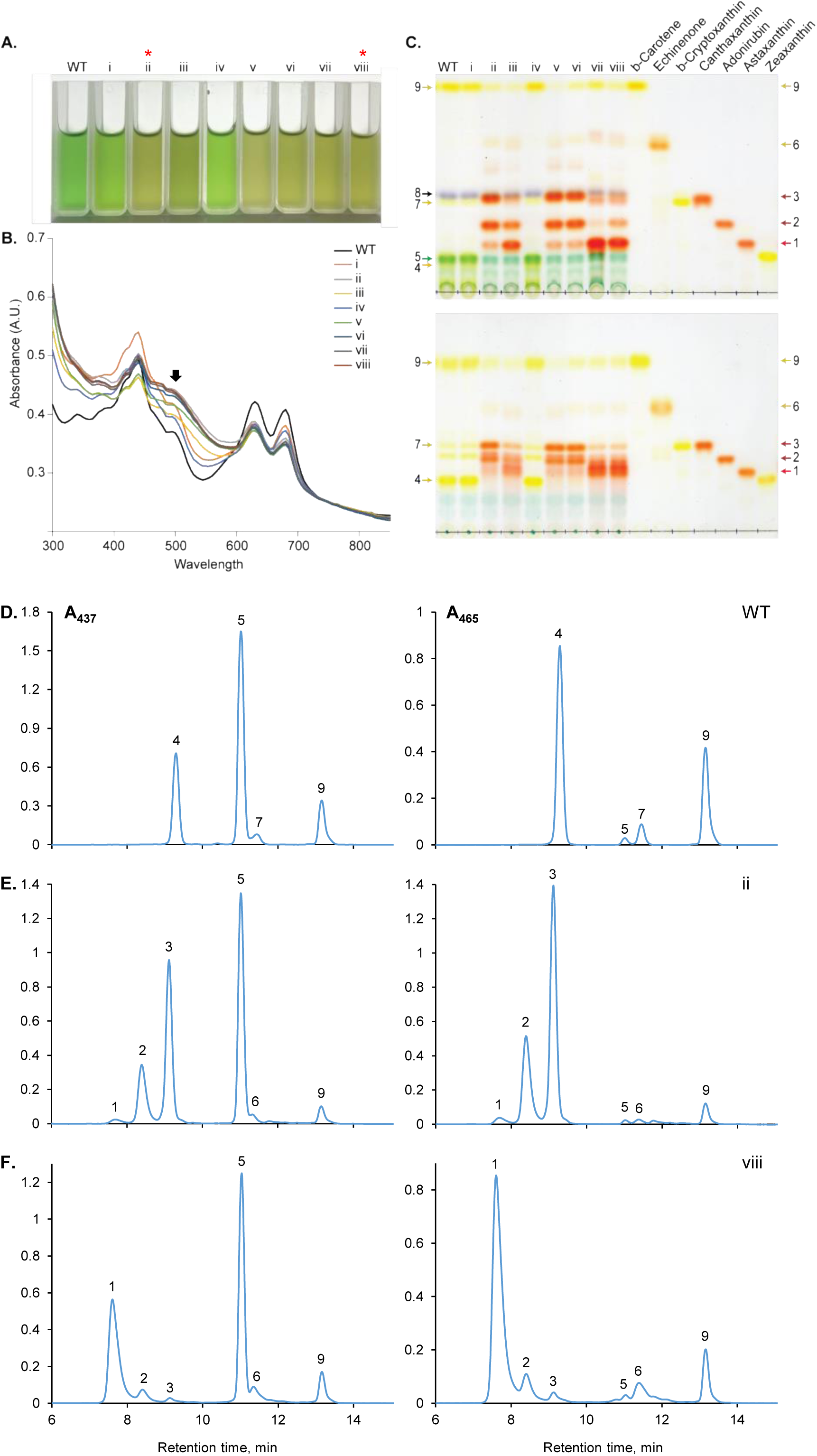
*C. merolae* 10D culture phenotypic changes and carotenoid profiles of transformants expressing different combinations of *Cr*BKT and *Cr*CHYB. **A** – Cuvettes containing 1 mL of *C. merolae* transformant culture for one representative of each confirmed plasmid transformation. **B** – absorbance spectra of cultures pictured above, shoulder of ketocarotenoid absorbance indicated with a black arrow. **C** – Acetone extract TLC of one confirmed representative *C. merolae* transformant for each indicated plasmid with carotenoid standards. Above – raw acetone extracts, below – saponified extracts. Arrows indicate 1 – astaxanthin, 2 – adonirubin, 3 – canthaxanthin, 4 – zeaxanthin, 5 – chlorophyll a, 6 – echinenone, 7 – β-cryptoxanthin, 8 – pheophytin a, 9 – *β*-carotene. HPLC profiles of carotenoids from parental *C. merolae* 10D (D), and transformants expressing *Cr*BKT– ii (E) or *Cr*BKT+*Cr*CHYB – viii (F).

TLC of acetone extracts then indicated the presence of orange-red pigments in transformants of each construct, which were absent from the parental or control strains expressing the CAT resistance alone or CAT and YFP alone (Figure 3C, plasmids i and iv). Transformants expressing variations of *Cr*BKT (plasmids ii, v, vi) were observed to accumulate canthaxanthin and adonirubin as major ketocarotenoids, with minor bands of astaxanthin (Figure 3C). The native carotenoid pathway contains hydroxylation activity to convert β-carotene into zeaxanthin (Figure 1). However, the accumulation of mostly canthaxanthin and adonirubin in *Cr*BKT expressing transformants indicates that the native CrtR activity does not outcompete the *Cr*BKT activity on β-carotene substrate and is not so highly active as to further hydroxylate these ketolated products. This is similar to the native CHYB activity in Chlamydomonas, which only creates significant titers of astaxanthin when overexpressed in the green alga as well (Amendola et al., 2023). Those transformants with co-expression of *Cr*CHYB with different fusion partners as well as *Cr*BKT (ii, vii, viii) exhibited minor bands of these two ketocarotenoids and astaxanthin as the major band in TLC (Figure 3C). Transformants of plasmid iii where *Cr*BKT and *Cr*CHYB are in a single fusion with each other, exhibited an intermediate phenotype, where astaxanthin was the major product, however, not as strong as with the two separate cassette expression in plasmids vii or viii. Patterns could be observed in non-saponified and saponified samples (Figure 3C, upper and lower panels, respectively). These patterns were true across individual transformants analyzed in a larger TLC with two representative transformants per plasmid construct is shown in Supplemental Figure S3.

Transformants were also subjected to flow cytometry analysis to determine the expression level of fusion reporter proteins, which confirmed the strength of expression for some constructs (Supplemental Figure S2). Plasmids iv and vi were shown to have fluorescence patterns distinct to those transformed with constructs harboring YFP fusions and those not harboring the YFP reporter (Supplemental Figure S2). Microscopy also confirmed localization in the chloroplast via YFP fluorescence (Supplemental Figure S2).

To determine the exact amounts of each carotenoid in the biomass, the parental strain and one transformant from plasmid ii (*Cr*BKT) and viii (*Cr*BKT+*Cr*CHYB) were subjected to pigment quantification by HPLC at 437 and 465 nm (Figure 3D-F; Table 1). Drastic differences in carotenoid profiles can be observed in the *Cr*BKT and *Cr*BKT+*Cr*CHYB expressing transformants. The *Cr*BKT expressing transformant exhibited a 33-61% reduction in β-carotene content accompanied by the disappearance of peaks #4 and 7 (zeaxanthin and *β*−cryptoxanthin, respectively) with the emergence of two predominant peaks #2 and 3 corresponding to adonirubin and canthaxanthin, respectively, small amounts of astaxanthin (peak #1). The *Cr*BKT+*Cr*CHYB expressing transformant consequently exhibited reductions in peaks #2 and 3 and significant increase in astaxanthin content (Figure 3D-F; Peak #1).

**Table 1.**
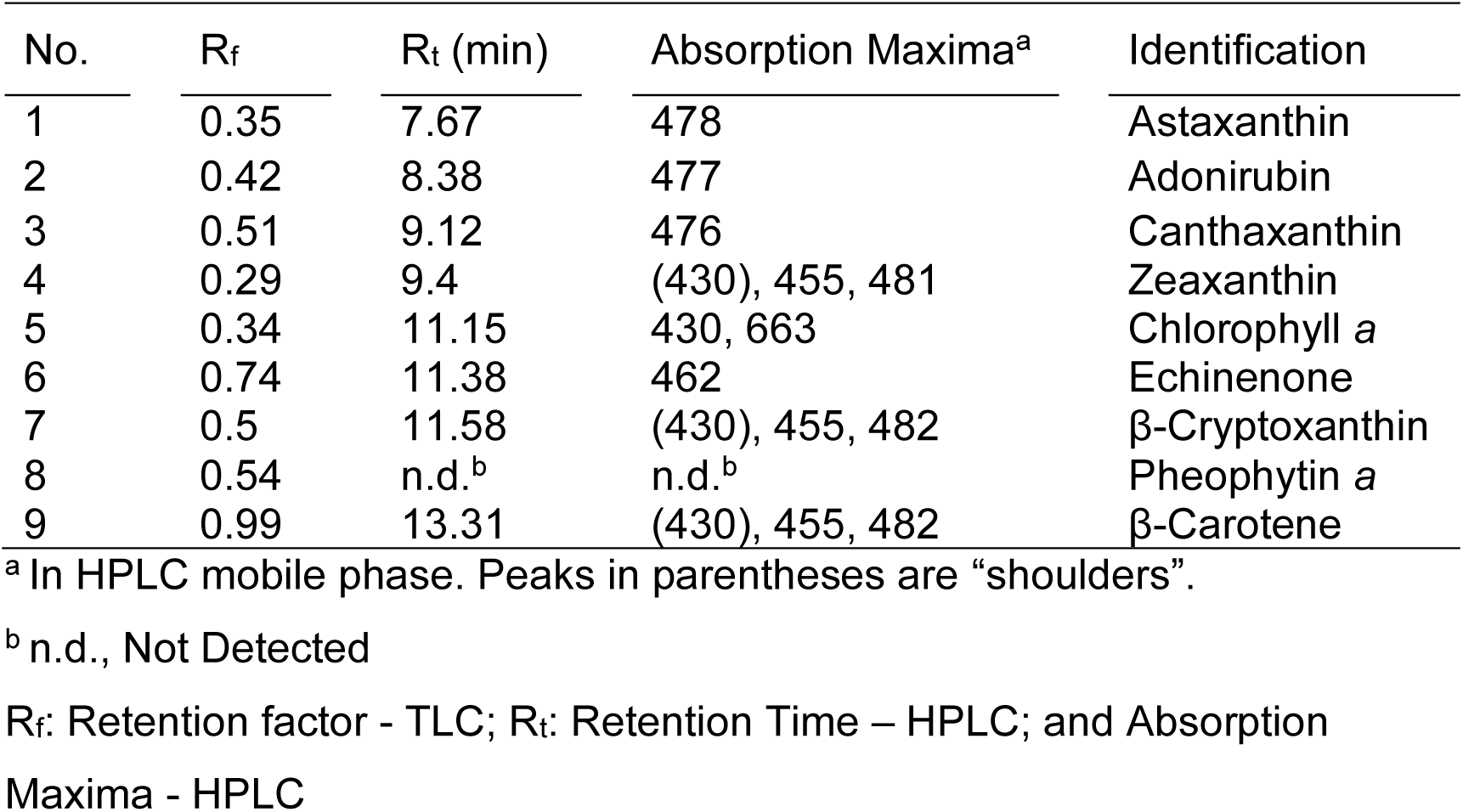
Identification of chlorophyll and carotenoid pigments in *C. merolae* WT and transformant lines: List of pigments detected and corresponding values of R_f_, R_t_ and absorption maxima are shown.

### Presence of ketocarotenoids improved total carotenoid content but slightly reduces growth rates of *C. merolae* 10D

Previous reports of ketocarotenoid biosynthesis in a green microalga indicated a global reduction of carotenoids and chlorophylls in transformants expressing *Cr*BKT but increased resistance to reactive oxygen species and high-light conditions (Amendola et al., 2023; Cazzaniga et al., 2022). It was unclear how ketocarotenoid presence would affect the photosystems of *C. merolae* here because these photosystems also contain phycocyanin as a light harvesting pigment and exhibit a natively minimal carotenoid profile lacking alpha carotenoids and terminal xanthophylls (Cunningham et al., 2007). The transformants and parental strain were subjected to a 12-day growth experiment in 20 mL culture tube (1.6 cm diameter) with 8 mL working volume that enable high-light penetration into the culture. Cultures were subjected to either 750 or 1172 μmol m^-2^ s^-1^ light intensity in a CO_2_ rich (3%) environment and sampled daily for growth metrics in addition to pigment quantification at the beginning, mid (L – log), and end of cultivation (S – stationary) (Figure 4A-H; Supplemental Data S5).

**Figure 4.**
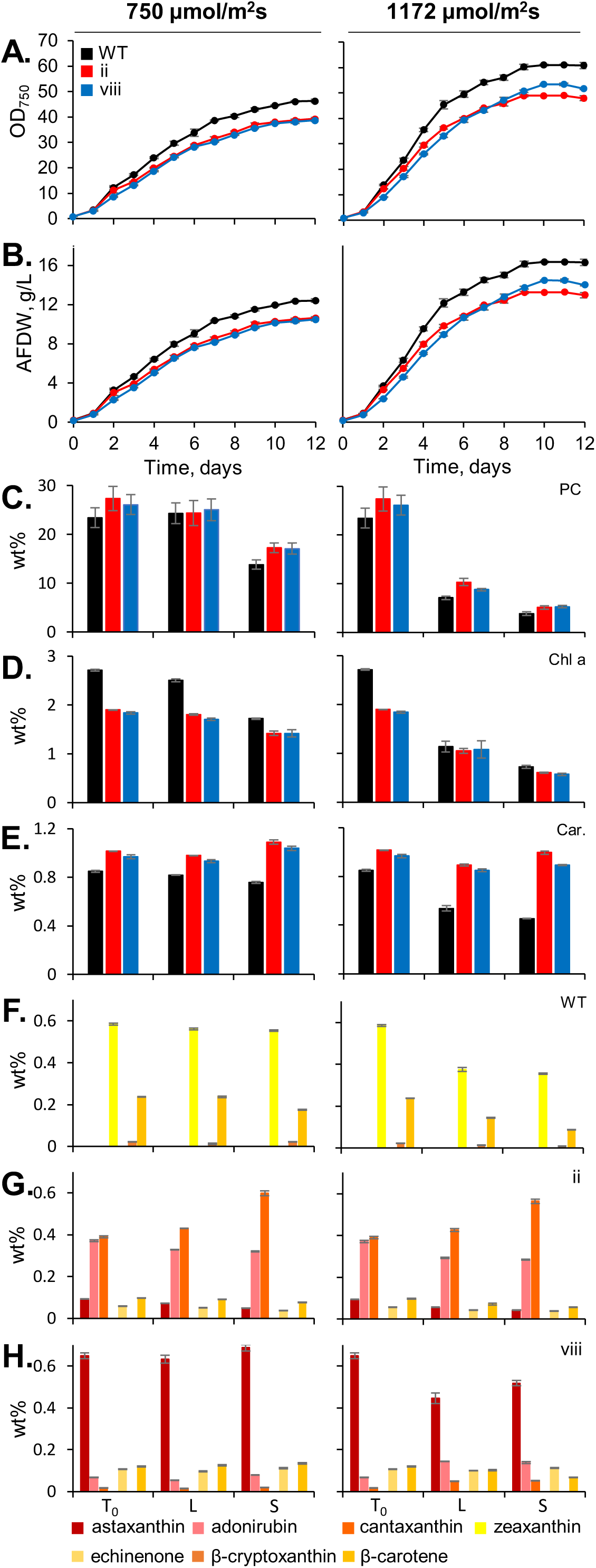
Growth behavior test and culture pigment profiles of parental (WT), *Cr*BKT, and *Cr*BKT+*Cr*CHYB transformants grown in 20 mL culture tubes under two light intensities. **(A)** Optical density (750 nm) and (B) ash-free cell dry weights (AFDW) were recorded throughout the 12-day cultivation. (A) phycocyanin, (B) Chlorophyll a and (C) total carotenoids were quantified at the start of cultivation, mid-log phase (d5), and stationary phase (d12), values are of the weight % of biomass. At each timepoint, the relative profiles of carotenoid species in each of the three cell lines ((F) WT, (G) *Cr*BKT – vii and (H) *Cr*BKT+*Cr*CHYB – viii) were also determined by HPLC and presented as weight % of the biomass.

In these optimized conditions, where light penetration into the thin culture tubes and CO_2_ are not limited, all cultures accumulated high rates of biomass over the 12-day period. *C. merolae* 10D achieved ∼12 g L^-1^ and ∼16 g L^-1^ in 750 and 1172 μmol m^-2^ s^-1^, respectively (Figure 4B). Growth behavior of both *Cr*BKT and *Cr*BKT+*Cr*CHYB transformants were ∼10 and 13-14 g L^-1^, respectively, in the two light conditions (Figure 4A and B). Phycocyanin content per cell was not significantly different between transformants and the parental strain in either illumination condition (Figure 4C). Total phycocyanin content was reduced in higher light conditions across all strains. Chlorophyll was overall lower in the higher light condition (Figure 4D), while total carotenoids were lower in the parental strain, but not in transformants (Figure 4E). Both types of pigments showed variation among the cell lines, with both ketocarotenoid accumulating strains exhibiting approximately 0.6-1.9 weight % chlorophyll a and approximately 0.9-1.1 weight % total carotenoid content (Figure 4 D and E).

The carotenoid profiles of each strain were unique, as shown in Figure 3 and Figure 4F-H, and trends observed in carotenoid species during the log phase were largely maintained in stationary phase for all cultures (Figure 4F-H). For the wild-type 10D, zeaxanthin was the most abundant carotenoid (0.35-0.58 weight %) with β-carotene as the second most abundant (0.09-0.24 weight %, Figure 4F). In the *Cr*BKT expressing strain, canthaxanthin was the most abundant carotenoid (0.39-0.60 weight %, with adonirubin (keto group on both terminal rings and single ring with hydroxyl group) the second most abundant (0.29-0.37 weight %, Figure 4G). In the *Cr*BKT+*Cr*CHYB expressing transformant, astaxanthin was the weight %, with adonirubin, canthaxanthin, echinenone, and β-carotene present but much less abundant (Figure 4H).

The results suggest that the total carotenoid per biomass in variable light conditions seems to be relatively constant despite reductions in overall other photosystem pigments in the ketocarotenoid producing transformants. Higher-light intensities reduced overall cellular phycocyanin contents, as expected based on previous reports of the behavior of this pigment in other organisms, where it is accumulated to assist photon capture in lower-light conditions (Chen et al., 2010). Similarly total chlorophyll reduction is also observed in higher light intensities, however, the red alga is unusual to what is observed in plants and green algae in that it does not have reduced overall carotenoids when ketocarotenoids are produced and in higher light (Cazzaniga et al., 2022; Perozeni et al., 2020). This could suggest that *C. merolae* is a promising chassis for tailored carotenoid production, especially considering it lacks a cell wall which enables simple carotenoid extraction. Concepts which aim to concomitantly acquire phycocyanin pigment and carotenoids from the same culture could use higher-light intensities to accumulate biomass as shown here and a period of lower light intensity before harvest to increase cellular phycocyanin yields, however, such tests were beyond the scope of this work.

### Modeling *C. merolae* growth in extreme environments

As a polyextremophile, *C. merolae* 10D can be grown in temperatures above cultivation norms for other algal species and in a very low pH to largely prevent contamination (Miyagishima and Tanaka, 2021). As the growth test performed in Figure 4 was performed in small culture tubes to ensure high-light penetration, we were curious how the parental and transformant strains would perform in larger culture volumes, where light penetration would become limiting. We grew the wild-type *C. merolae* 10D, *Cr*BKT (ii), and *Cr*BKT+*Cr*CHYB (vIII) strain in 400 mL cultures using a suite of photobioreactors to tightly control environmental parameters while tracking growth (Figure 5). Cell lines were grown at 42 °C with constant 1500 µmol photons m^-2^ s^-1^ illumination or 12:12 day:night light cycling to represent controlled bioreactor cultivations in optimal conditions. In addition, we used collected weather data generated on the mid-Red Sea coast (Supplemental Data S6) and in Mesa Arizona (Supplemental Data S7) to generate 8-day cultivation programs representing one month of each season in these locales. The summer months in both geographies exhibit high temperatures, with Mesa having higher midday temperatures and greater fluctuations between day and night (Figure 5A-D, right panels).

**Figure 5.**
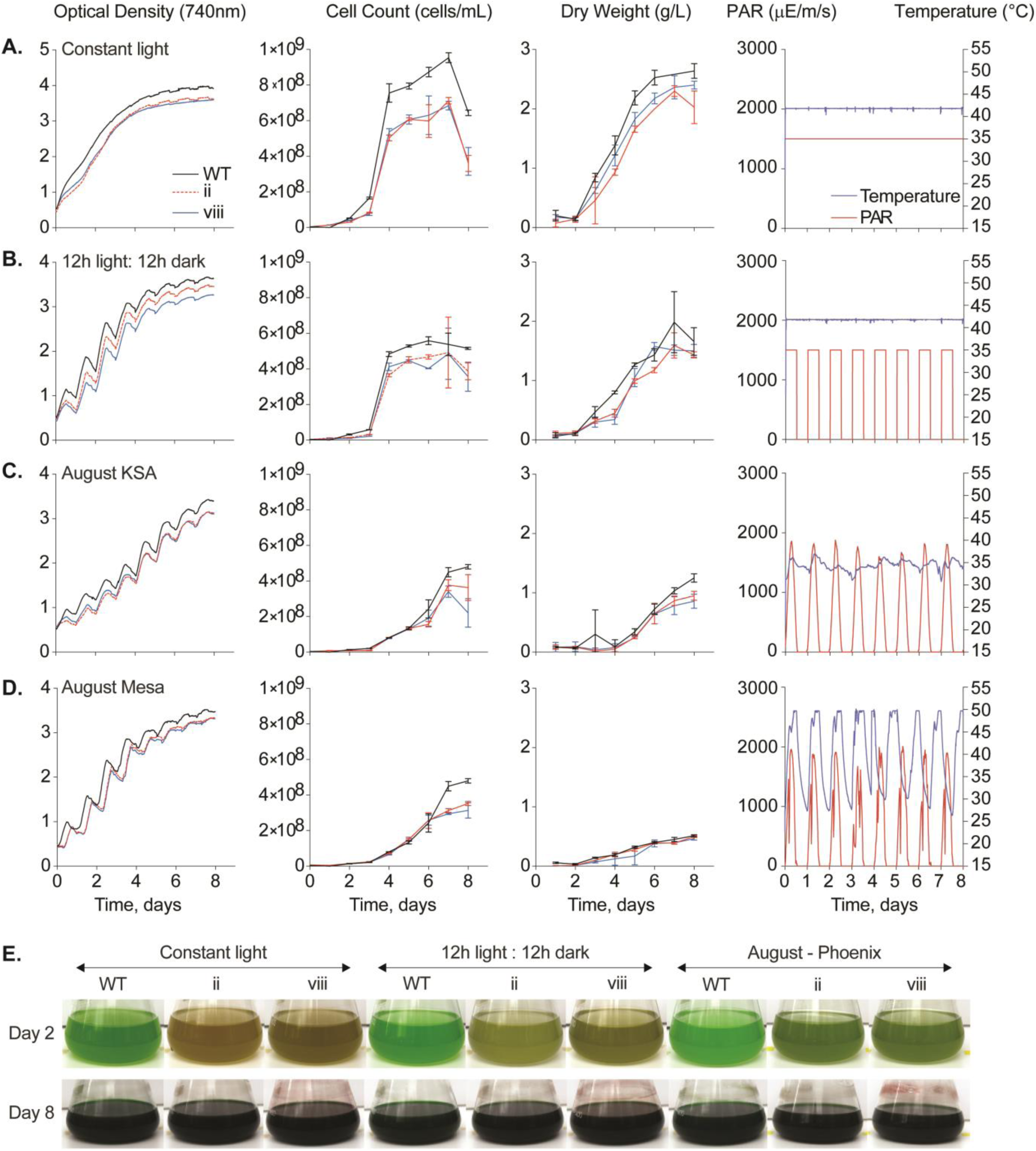
Comparative bioreactor growth tests of parental *C. merolae* 10D, *Cr*BKT, and *Cr*BKT+*Cr*CHYB transformants in various conditions. The three cell lines were cultivated in (A.) constant 1500 µE illumination and with (B.) 12:12 hour day:night cycling at 42 °C as well as simulated environmental conditions from recorded weather data for the month of August in the (C.) mid-Red Sea coast (KSA) and (D.) Mesa Arizona. Optical density (OD 740 nm), cell density (cells/mL), and dry biomass (g L^-1^ culture) are indicated beside the light and temperature profiles used in each bioreactor. One cultivation of three biological replicates is shown. Below (E.), culture flask pictures at day 2 and 8 of the cultivation showing phenotypic differences in ketocarotenoid accumulating transformants.

In all bioreactor conditions, the ketocarotenoid producing transformants exhibited slightly lower optical and cell densities, as well as biomass compared to their parental strain (Figure 5A-D). Both transformants performed similarly, suggesting that the presence of ketocarotenoids at all, rather than a specific type, caused this growth behavior difference. In continuous illumination, the 400 mL cultures achieved ∼2.5 g L^-1^ biomass in 6 d, while the ketocarotenoid transformants accumulated ∼2.2 g L^-1^ (Figure 5A-D). Overall cell densities exhibited similar amounts in both geographies, with mid-Red Sea coast having slightly higher biomass accumulated than in Mesa (Figure 5A-D). The higher temperatures observed in Arizona summer exceeded the capacity of the bioreactor (+50 °C), temperatures which would likely be detrimental to many algal species in culture (Figure 5). Nevertheless, it was still possible to grow both transformed and parental *C. merolae* in this condition where they accumulated the ketocarotenoid products (Figure 5E, pictures). All data for phycocyanin, chlorophyll, and carotenoid accumulation can be found in Supplemental Figure S4.

### The value of *C. merolae* 10D as a host for engineered carotenoid biosynthesis

The Cyanidiales are polyextremophilic red algae which have emerged in recent years as interesting alternatives to other algal systems (Lang et al., 2020). Within this class are several species that are found in acidic hot springs and thrive between pH 0.5-5 and temperatures from 35-56 °C. These growth conditions set the Cyanidiales apart from other algae in that few organisms can grow in such conditions and contamination at scale can be largely prevented. *Galdieria sulphuriana* is another species within this Class that has been shown to be capable of rapidly becoming the dominant organism when grown directly in acidified municipal effluent (Henkanatte-Gedera et al., 2017, 2015). *C. merolae* 10D is a obligate phototroph and can only consume CO_2_ as a carbon source (Miyagishima and Tanaka, 2021). It is also tolerant to very high levels of CO_2_ gas, ammonium concentration in its medium, and high temperatures (Minoda et al., 2004; Miyagishima and Tanaka, 2021). These features potentially mean *C. merolae* 10D could be coupled to post-treatment high-strength wastewater polishing and industrial CO_2_ emissions sources in extreme conditions such as those in desert environments modelled here.

*C. merolae* 10D is also interesting for biotechnological applications owing to its lack of cell wall and range of native natural products which can be rapidly separated in various phases of extraction. The cell itself contains a small lipid fraction, starch, and β-glucan in addition to ∼50% protein content (Miyagishima and Tanaka, 2021). *C. merolae* accumulates the photopigment phycocyanin which is water-soluble and more thermal stable than that currently used in industry produced by *Arthospira platensis* (Rahman et al., 2017). The parental strain also accumulates large fractions of zeaxanthin and β-carotene which are both valuable hydrophobic pigments (Figures 3 & 4). The cell, therefore, is a natural candidate for biorefinery concepts, as PC and soluble proteins can be readily extracted from cell-wall-less biomass and carotenoid pigments isolated from the residual insoluble fraction. Separate fractionation of starch and β-glucans may also be possible with appropriate bioprocess designs. This concept is illustrated in Figure 6.

**Figure 6.**
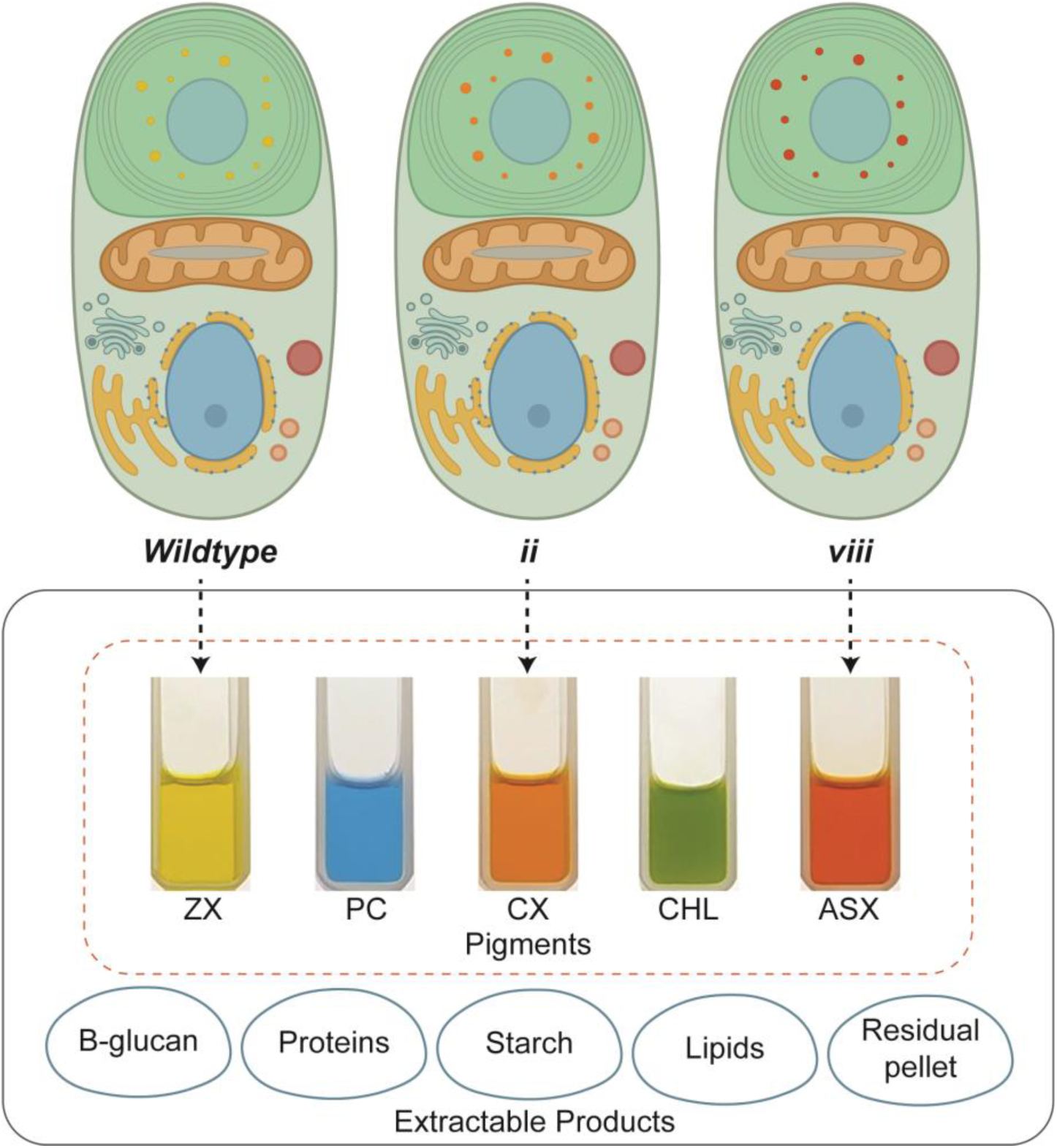
Extractable products from wild-type and engineered *C. merolae* 10D. The schematic displays the extractable products that can be obtained from *C. merolae* 10D cells through various extraction phases. The dotted arrows indicate the carotenoid fractions that can be extracted from the corresponding cell lines: WT, ii (*Cr*BKT), and viii (*Cr*BKT+*Cr*CHYB). The pigment fractions are named based on the predominant carotenoid present in the extract: ZX (zeaxanthin), CX (canthaxanthin), and ASX (astaxanthin). Additionally, phycocyanin (PC) and chlorophyll a (CHL) are present in all lines. Pigments were extracted as described in M&M and ∼1 mL of each was photographed in 3 mL cuvettes.

The capacity for engineering ketocarotenoid biosynthesis expands the product range which can be achieved from this easy-to-handle organism, with *Cr*BKT expression producing canthaxanthin and the combination of *Cr*BKT+*Cr*CHYB astaxanthin. Our results indicate that despite a subtle reduction in overall growth rates when cells produce ketocarotenoids (Figure 4 and 5), they are still amenable to cultivation in extreme conditions and do not reduce their overall carotenoid contents, even in high light conditions. Future optimization of cultivation parameters can tease-apart the best light and temperature regimes to promote biomass accumulation and increase cellular classes of photopigments in the engineered cells. Our work indicates that *C. merolae* 10D could be cultivated outdoors, even in some of the hottest urban environments in the world during summer months, but life-cycle analysis would be required to determine whether the CAPEX required to build a controlled bioreactor with constant illumination would be more beneficial than simply using outdoor environmental conditions *in situ*. This is also encouraged by our recent finding that *C. merolae* can be adapted to be grown in acidified sea water salinities, further expanding its possible range of geographical application (Hirooka et al., 2020; Villegas et al., 2023). Indeed, each implementation of such a cultivation would require individual case-considerations. The thermal extreme tolerance of *C. merolae* 10D and its engineered derivatives at least suggests that cooling will not be needed if bioreactors are placed outdoors. Waste-heat may be used to optimize culture conditions, especially during colder seasons, as this is energetically less challenging to engineer into a cultivation apparatus than cooling in these extreme environments.

## 4. Conclusions and Outlook

Here, we show the power of *in silico* design and *de novo* construction of transgene expression constructs in an emerging host microalga. We used these molecular tools to rapidly demonstrate the production of non-native ketocarotenoids in the polyextremohilic red microalga which has emerged in recent years as a promising alternative to other green algal hosts. This work represents the first demonstration of carotenoid metabolic engineering by recombinant technologies in any red alga. The lack of impact alternative carotenoid production had on soluble phycocyanin contents adds interesting value to an already specialized algal biomass as these products can be separately extracted as soluble and insoluble fractions from the biomass. The wild-type strain is already a source of zeaxanthin, and our findings indicate it is possible to tailor this host into a production vehicle for either canthaxanthin or astaxanthin without contaminating alpha carotenoids. Given each of these carotenoids has a value of their own, parallel cell lines could be used to generate multiple products from the same algal cultivation infrastructure. Adaptability to saline conditions, high temperature tolerance, and the capacity for growth on high strength waste-waters also encourage the potential value economics of *C. merolae* bio-production processes. Given the relative ease of transgene integration into the nuclear genome of this algae and high expression rates, it will likely rapidly become a host cell for a range of photosynthetic engineering concepts.

## Supporting information

Supplemental Figure S1 Primer anealing locations

Supplemental Figure S2 YFP fluor analysis

Supplemental Figure S3 TLC unsap and sap

Supplemental Figure S4 algem pigment analysis

Supplemental Data S1 - Sequence information

Supplemental Data S2 - Primers and PCR assay

Supplemental Data S3 - Carotenoid standard curves

Supplemental Data S4 - plasmids annotated

Supplemental Data S5 - Indoor culture tube experiments

Supplemental Data S6 - Thuwal growth tests and data

Supplemental Data S7 - Mesa growth tests and data

## Abbreviations

CDW: cell dry weight
YFP: mVenus yellow fluorescent protein
*Cr*BKT: *Chlamydomonas reinhardtii* β-carotene ketolase
*Cr*CHYB: *C. reinhardtii* β-carotene hydroxylase
CAT: Chloramphenicol transferase
CTP: chloroplast targeting peptide
AFDW: ash free dry weight
PC: phycocyanin
HR: homologous recombination
TLC: thin layer chromatography
HPLC: high-performance liquid chromatography

## Acknowledgements

KJL acknowledges baseline research funding provided by King Abdullah University of Science & Technology. KAUST team is grateful to Paulo C. Aurelio of KAUST Core Labs Lab Equipment Maintenance (LEM) team for install and maintenance of the Algem photobioreactors and flow cytometer. PJL acknowledges financial support from Xylem, Inc. and ASU Lightworks; and would also like to acknowledge Keirsten Allen for her technical support. The authors wish to express their gratitude to Dr. Martha Stark for her invaluable assistance with the transformation protocol. The authors thank Dr. Sebastian Overmans for creating Algem bioreactor profiles from provided data. The authors thank Dr. Thomas Baier for invaluable discussions around the use of CHYB in *C. reinhardtii* and sharing these insights with us.

## Supplemental Figures

**Figure S1: Schematic map of primer annealing sites for the targeted integration site at the 184C-185C locus, on the transformation plasmids, and the RuBisCO large subunit locus**. Additional information on the primers used can be found in supplemental Table S2, including the PCR assay conditions. The primer sets were utilized for the following purposes: (1) to determine the presence or absence of each transgene (*Cr*CHYB, YFP, *Cr*BKT, and CAT), (2) to confirm the integration of DNA insert (using 209F/2776R for arms and M2F/D184R for integration, anchored outside of insert), (3) to check for the presence or absence of plasmid DNA (using EpiF/R for episomal), and (4) to verify the unialgal status of the cultures (using universal RbcL R/F for RuBisCO LSU followed by *Hae*III R.E. digestion). Red and blue triangles indicate the forward and reverse primers, respectively, and lines (dotted or solid) connect the primer sets.

**Figure S2: Evaluation of YFP transformants via flow cytometer and epifluorescence microscopy**. (A.) Histograms of the forward scatter (FSC), side scatter (SSC), chlorophyll fluorescence (Chl; 692/40 nm), and YFP fluorescence (YFP; 575/25 nm) of wild type and YFP transformant cells. (B.) Brightfield and epifluorescent images of transformant cells expressing YFP to verify chloroplast localization. Representative image shown using transgenic line iv. Brightfield image in left top corner, pigment and YFP fluorescence shown in top right and bottom left corner, respectively. Overlay of all three shown in bottom right corner.

**Figure S3: Carotenoid profiles of WT and transformants expressing different combinations of *Cr*BKT and *Cr*CHYB**. Acetone extract TLC of two *C. merolae* WT and confirmed representative transformants for each indicated plasmid. The raw acetone extracts are displayed above, and the saponified extracts are shown below. Arrows indicate the following carotenoids: 1 - astaxanthin, 2 - adinorubin, 3 - canthaxanthin, 4 - zeaxanthin, 5 - chlorophyll a, 6 - echinenone, 7 - β-cryptoxanthin, 8 - pheophytin, 9 - β-carotene.

**Figure S4**: **Pigment analysis results of an Algem growth experiment conducted under Mesa and Thuwal simulated environmental conditions**. Chlorophyll (A), total carotenoids (B), and Phycocyanin (C) were extracted from the WT, ii (*Cr*BKT), and viii (*Cr*BKT+*Cr*CHYB) lines on day 4 and day 8 of cultivation in Algem® photobioreactors. The experiment was carried out under four different growth conditions: Constant (24h) and diurnal (12h:12h) light conditions at 1500 μmol m^-2^ s^-1^ and 42 °C; and two simulated environmental conditions for the month of August in Thuwal (Saudi Arabia) and Mesa (Arizona). The averages of biological triplicates are displayed.

## Supplemental data captions

**Supplemental Data S1 Sequences and sources used to construct transformation plasmids**. Table includes abbreviated names, function, gene name and ID, sequence with indicated modifications and size, along with source (organism and references).

**Supplemental Data S2 – Primer and PCR assay information**. Table includes comprehensive information on the primers used for this study to screen and monitor transformants. Target templates/genes, primer abbreviations, sequences along with Tm’s, product sizes, and PCR assay conditions are provided.

**Supplemental Data S3 – HPLC standard quantifications** Calculations for carotenoid standard curves. Standard curves like the ones shown here were used to quantify carotenoids during experimentation. Standards were also used to confirm bands in TLC (Figure 3) and peaks (retention times and absorption spectra) in HPLC chomatograms.

**Supplemental Data S4 – Annotated sequences of all plasmids used in this work**. This file can be opened with any plasmid editor software to see plasmid sequences and annotation maps.

**Supplemental Data S5 – Indoor culture tube experiments**. Consolidated data set for the indoor 20 mL culture tube experiment. Data was used to produce Figure 4.

**Supplemental Data S6 – Thuwal conditions growth tests and data**. Consolidated data set for the Algem photobioreactor simulating environmental conditions for Thuwal, Saudi Arabia. Data was used to produce Figure 5C.

**Supplemental Data S7 – Mesa conditions growth tests and data**. Consolidated data set for the Algem photobioreactor simulating environmental conditions for Mesa, Arizona. Data was used to produce Figure 5D.

