## Supplementary figures and images for "Engineered ketocarotenoid biosynthesis in the polyextremophilic red microalga *Cyanidioschyzon merolae* 10D"

### Supplemental Figure S1 Primer anealing locations

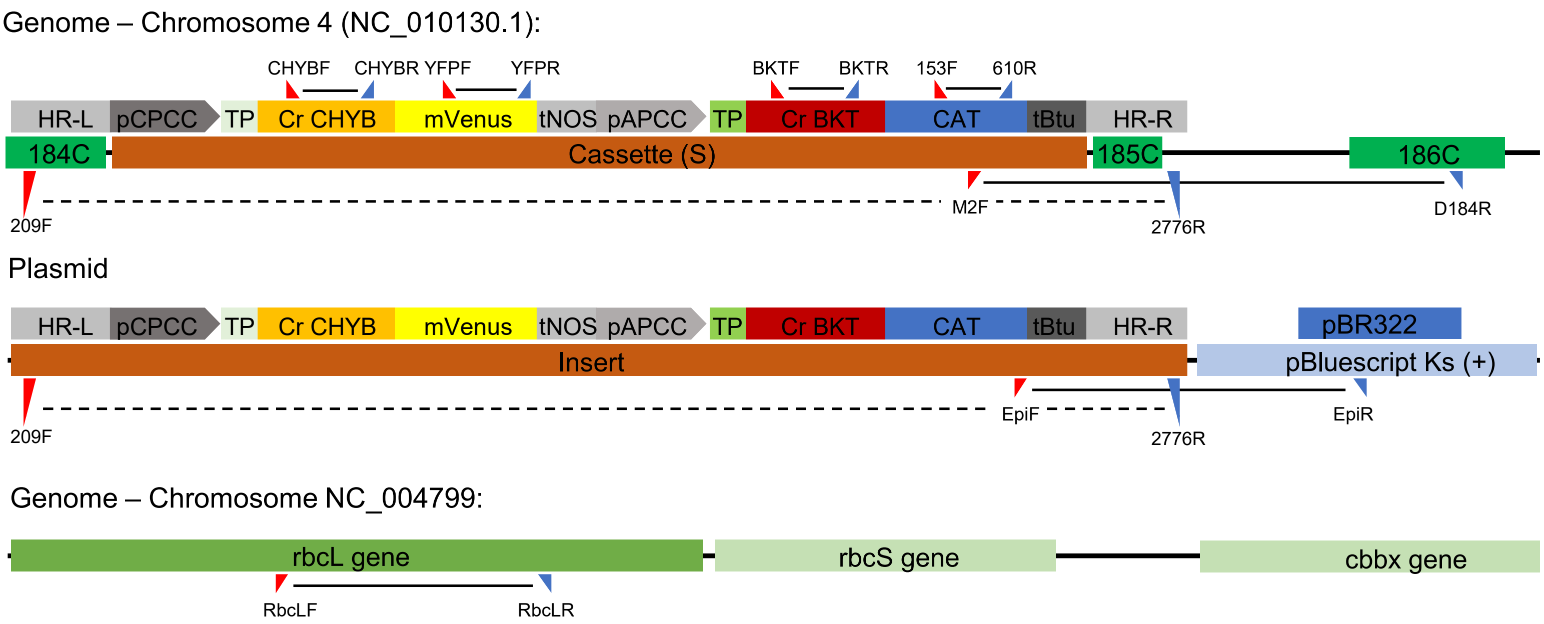

### Supplemental Figure S2 YFP fluor analysis

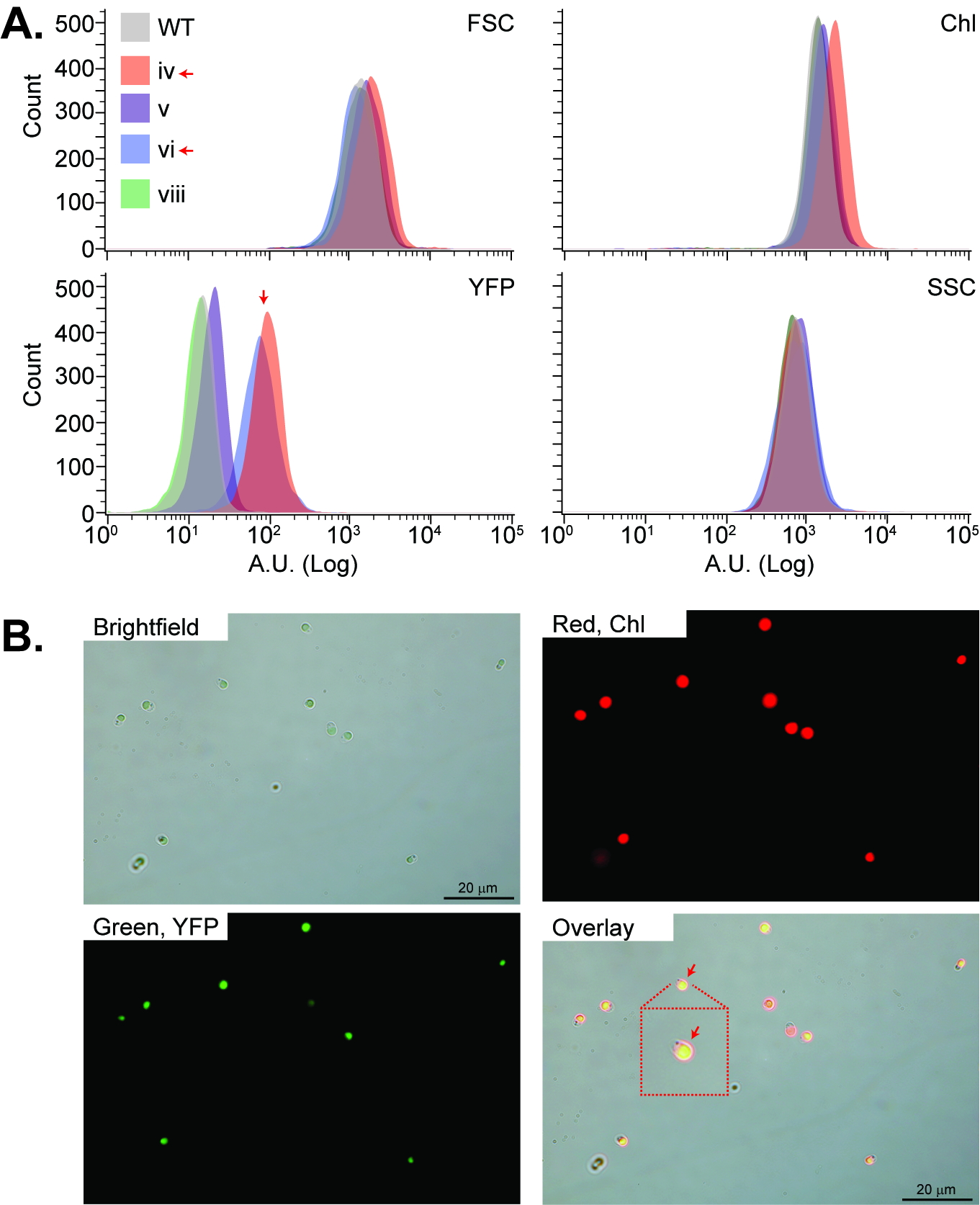

### Supplemental Figure S3 TLC unsap and sap

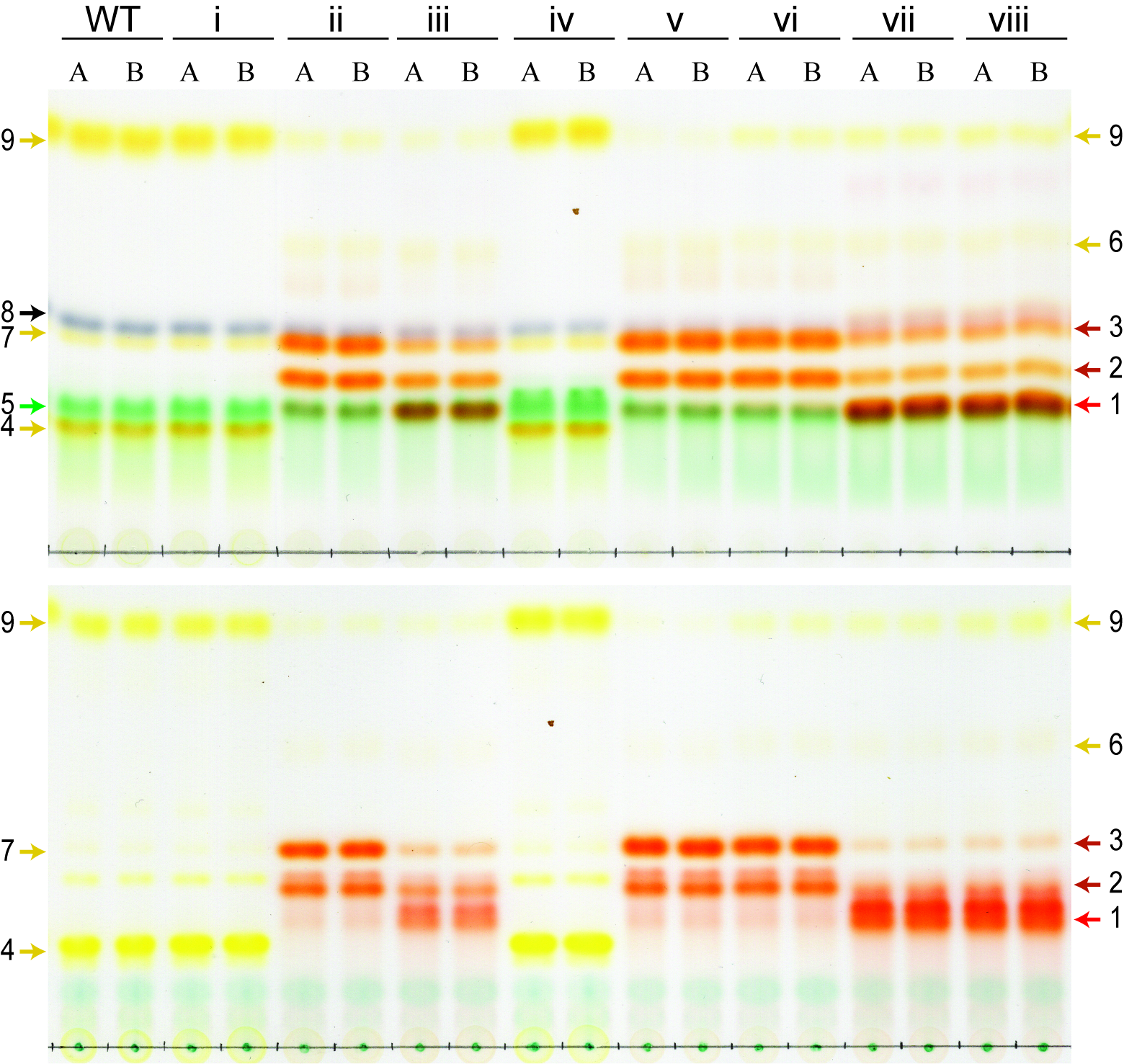

### Supplemental Figure S4 algem pigment analysis

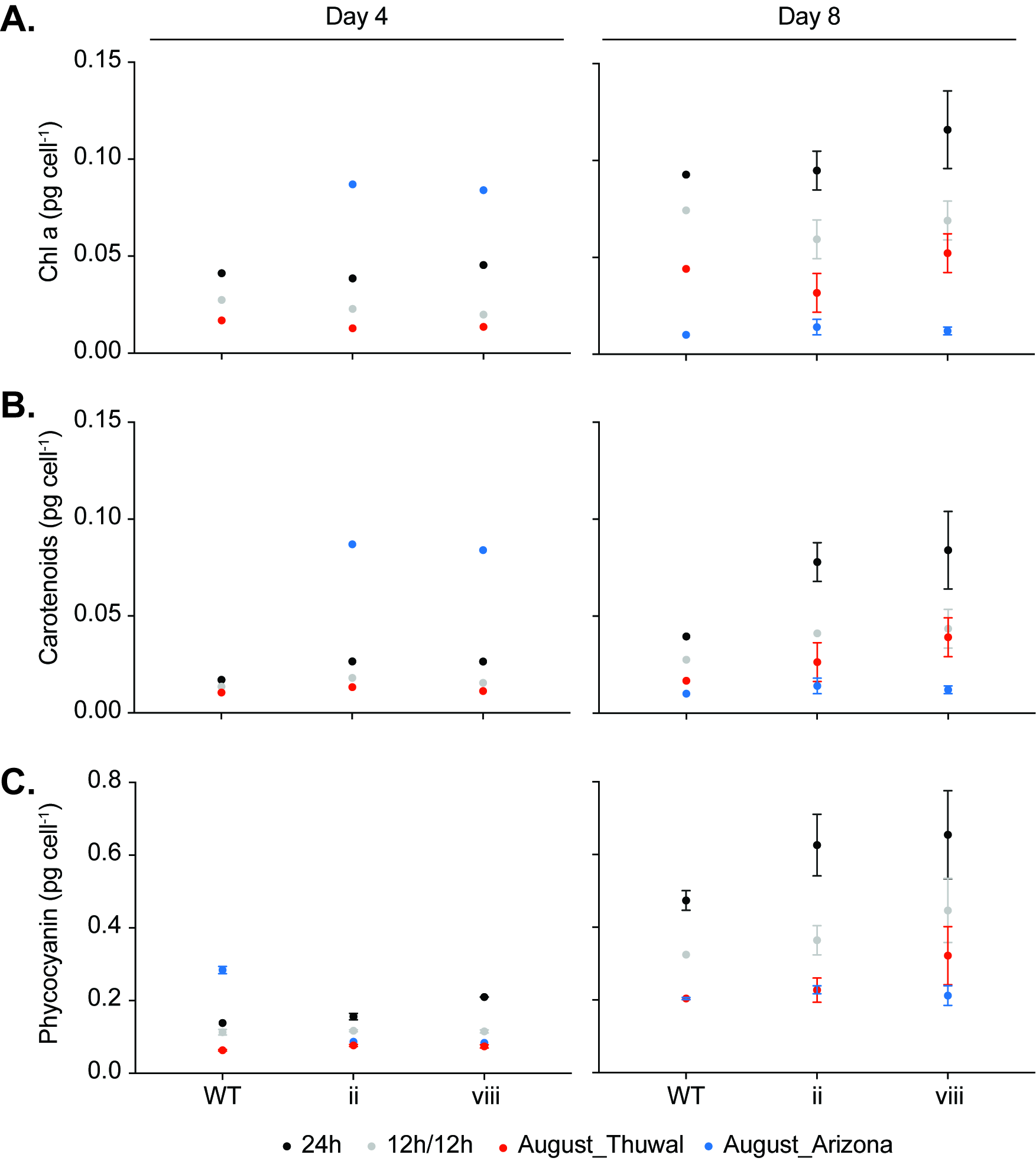
